# RUCova: Removal of Unwanted Covariance in mass cytometry data

**DOI:** 10.1101/2024.05.24.595717

**Authors:** Rosario Astaburuaga-García, Thomas Sell, Samet Mutlu, Anja Sieber, Kirsten Lauber, Nils Blüthgen

## Abstract

High dimensional mass cytometry is confounded by unwanted covariance due to variations in cell size and staining efficiency, making analysis and interpretation challenging.

We present RUCova, a novel method designed to address confounding factors in mass cytometry data. RUCova removes unwanted covariance using multivariate linear regression on Surrogates of Unwanted Covariance (SUCs), and Principal Component Analysis (PCA). We exemplify the use of RUCova and show that it effectively removes unwanted covariance while preserving genuine biological signals. Our results demonstrate the efficacy of RUCova in elucidating complex data patterns, facilitating the identification of activated signalling pathways, and improving the classification of important cell populations. By providing a robust framework for data normalization and interpretation, RUCova enhances the accuracy and reliability of mass cytometry analyses, contributing to advancements in our understanding of cellular biology and disease mechanisms. The R package is available on https://github.com/molsysbio/RUCova. Detailed documentation, data, and the code required to reproduce the results are available on https://doi.org/10.5281/zenodo.10913464. Supplementary material: Available at bioRxiv.

## Introduction

Mass cytometry allows the simultaneous quantification of numerous cellular markers in individual cells and across multiple samples. It is used widely in immunology research to quantify surface proteins and classify immune cells [Spitzer and Nolan, 2016; Bendall et al., 2011; Horowitz et al., 2013; Giesen et al., 2014; Georg et al., 2022]. Mass cytometry is also increasingly used to study intracellular signalling pathways by measuring phospho-protein abundance, providing insights into diverse cellular processes such as the differentiation pathways of colorectal cancer [Brandt et al., 2019; Sell et al., 2023], organoid heterogeneity [Sufi et al., 2021], acute myeloid leukaemia [Han et al., 2015] and prediction of drug sensitivity in breast cancer [Tognetti et al., 2021]. While the distributions of surface proteins typically show a bimodal pattern, those of intracellular signalling markers show a unimodal distribution with rather small quantitative shifts in response to perturbations. These distributions are affected by both biological and technical variability. Biological variability arises from inherent differences between individual cells, including variations in cell state, type and size, whereas technical variability arises from experimental procedures and instrumentation, such as heterogeneous staining efficiency. While some biological variability is essential, unwanted variability, such as that caused by differences in cell size, carries the risk of confounding the data. This unwanted covariance can obscure the detection of small cell populations and prevent accurate comparisons between different experimental conditions, cell lines and cell states.

In recent years, a class of methods called Remove Unwanted Variation (RUV) has been developed to address this problem, primarily variation coming from batch effects. These methods have been successfully applied to various high-throughput data types, including microarrays [Gagnon-Bartsch and Speed, 2011], RNA-seq (RUV by Risso et al. [2014]), Nanostring nCounter gene expression (RUV-III by Molania et al. [2019]), single-cell RNA-seq (scMerge by Lin et al. [2019]), and mass cytometry (CytofRUV by Trussart et al. [2020]). While differences between batches are a significant source of unwanted variability, single-cell mass cytometry datasets can exhibit considerable covariance within a single batch due to uncorrected heterogeneity in cell size and staining efficiency. Mass cytometry builds upon flow cytometry by increasing the number of measurable markers, as the mass spectrum is more specific than the fluorescence spectrum, which suffers from spectral overlap. In flow cytometry, marker abundance can be normalized by cell size using the Forward Scatter (FSC) parameter, which is proportional to the cell’s relative size. However, mass cytometry lacks an intrinsic parameter that directly serves as a proxy for cell size. Conventional normalisation methods, such as those used in single-cell RNA sequencing, are impractical due to the absence of total protein content information in mass cytometry data. Previous attempts to normalize cell volume using Ruthenium isotopes [Rapsomaniki et al., 2018] faced challenges with complexity and assumptions about marker-cell volume relationships. To address these limitations, we introduce RUCova, a novel approach utilising linear modeling based on Surrogates of Unwanted Covariance (SUCs). By incorporating factors like mean DNA, mean barcoding isotopes, pan Akt, and total ERK, RUCova effectively removes technical artifacts and enhances the accuracy of mass cytometry analyses. Our study demonstrates the utility of RUCova in revealing complex data patterns, identifying activated signalling pathways, and improving the classification of important cell populations like apoptotic cells. In this paper, we showcase the unique advantages of RUCova in advancing our understanding of cellular biology and disease mechanisms.

### The RUCova method

RUCova comprises two major steps. First, it fits a multivariate model for each measured marker (*m*) across cells (*i*) from samples (*j*_*i*_) with respect to the surrogates of unwanted covariance (SUC)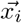. Second, it eliminates such dependency by assigning the residuals *ϵ* of the model as the new modified expression of the marker. The fit can be expressed as:

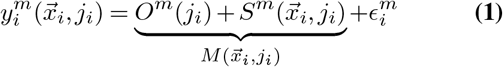

where 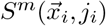 describes the slope of the fit and *O*^*m*^(*j*_*i*_) the intercept or offset. The predictors are the surrogates of unwanted covariance (SUC) 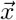. which can be either specific markers as proxies of the confounding factors or the principal components (PCs) derived from a Principal Component Analysis (PCA) performed on such markers.

RUCova offers 3 different uni– or multi-variate linear models to describe the relationship between marker expression and SUC: (1)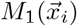: simple, (2)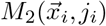: offset, and (3) 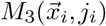: interaction (Fig. 1).

**Fig. 1.**
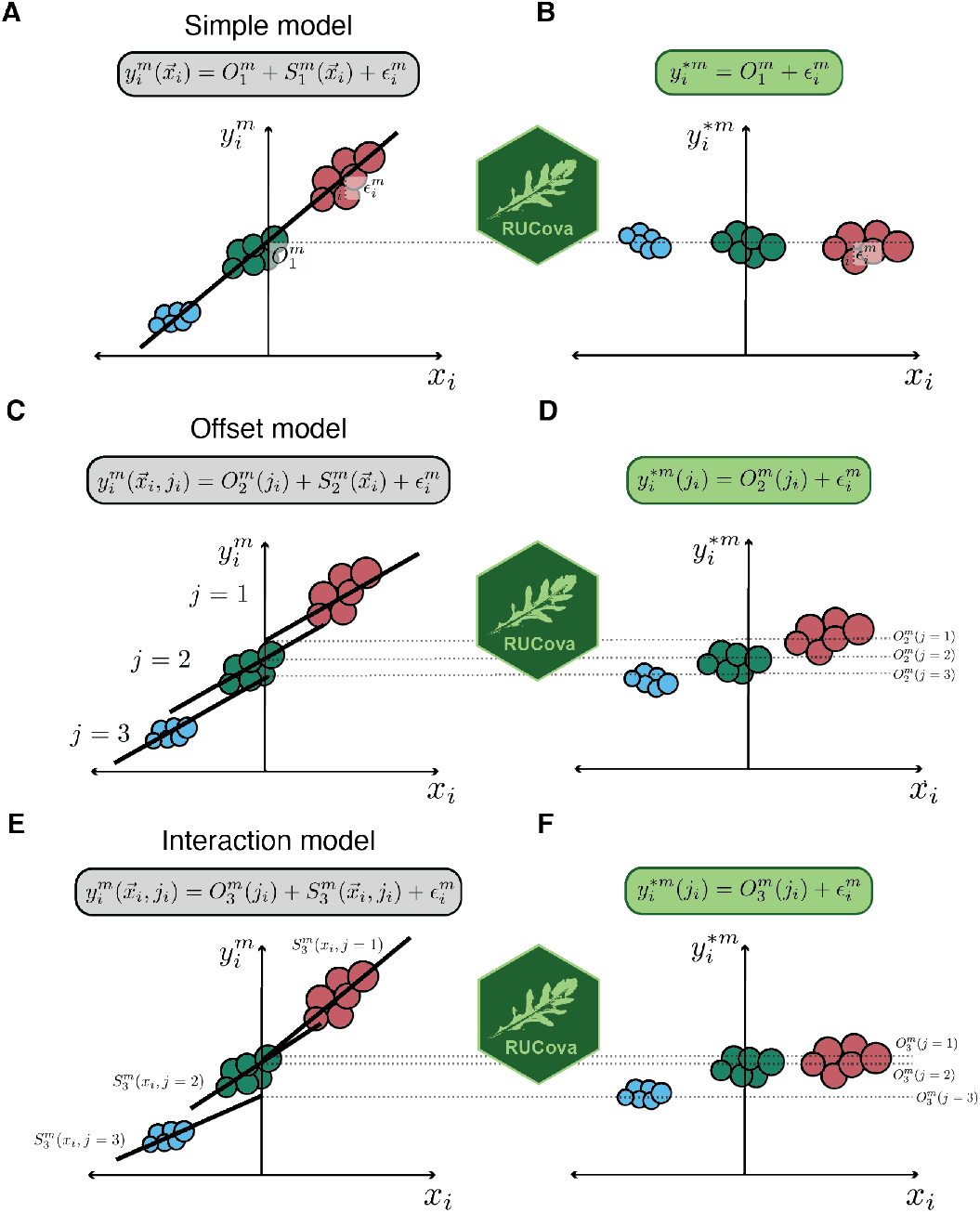
Illustration of the RUCova method and its three different models. **A, C, E)** Original expression 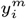 of a marker *m* before RUCova as a function of a centred expression of a SUC or PC. Illustrative regression line and equation corresponding to each model. **B, D, F)** Modified expression 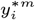 of a marker *m* after applying RUCova. **A, B)** Simple model: one fit across the input data set with intercept 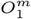 and residuals *ϵ*_*i*_. **C, D)** Offset model: one slope 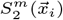 for the whole input data set and different intercepts 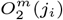 between samples. **E, F)** Interaction model: one fit per sample *j* with intercepts 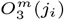 and slope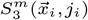. **D, F)** Keeping the offset *O*^*m*^(*j*_*i*_) between samples *j*.

1. **Simple model (Fig. 1A**,**B):** Consists of one fit per measured marker 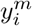 across the entire data set.

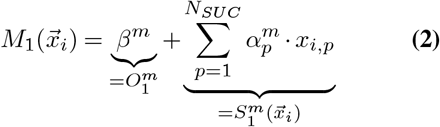

where 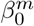 is the intercept and 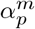 is the slope coeffi-cient for each predictor or SUC *p*.
2. **Offset model (Fig. 1C**,**D):** Consists of one fit per measured marker 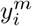 and sample *j*_*i*_. The fits for the sam-ples share the same slope, while differing in the intercept (offset term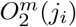).

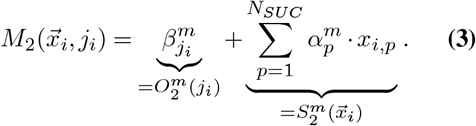
3. **Interaction model (Fig. 1E**,**F):** Consists of one fit per measured marker 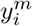 and sample *j*_*i*_. The fits for the samples can have different slopes (interaction term 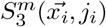) and intercepts (offset term 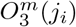).

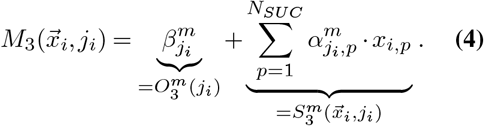

Samples *j*_*i*_ can be either different cell lines, perturbations, conditions, *metacells* (clusters), or even batches. By taking the zero-centered distributions of the SUCs 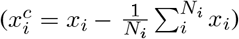, the mean values of the markers *m* across all cells are kept after applying RUCova (Fig. 1). If a more conservative approach is desired where the fold changes between samples should be kept, each SUC should be centred per sample (Fig. S1). Similarly, when using PCs as the predictive variables, SUCs can be z-score normalised by the sample before performing PCA.

The RUCova method eliminates the dependency of each measured marker on the SUCs by computing the model’s residuals and the intercept as the revised expression for each marker 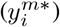. The offset – intercept – term *O*^*m*^ for different samples can be wanted or unwanted. For the first case, the new, modified abundance of the marker after applying RUCova is independent of 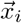 and can be expressed as:

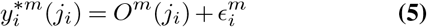

More information about the RUCova model, cell cultures and mass cytometry measurements can be found in the Supplementary Material.

## Results

### 1. Mass cytometry data is confounded by multiple factors

Mass cytometry enables the quantification of protein and phospho-protein abundance in single cells using antibodies conjugated with metal isotopes, facilitating the investigation of intracellular signals that determine the state and response to treatments. However, challenges such as heterogeneous cell volume and labelling efficiency confound the data, leading to spurious correlations between markers and hindering comparisons between cell lines, perturbations, and cell states. To address this, we developed RUCova, an R package designed to remove unwanted covariance in mass cytometry data.

To illustrate the need and benefit of using the RUCova method, we chose a mass cytometry data set with 8 different Head-and-Neck Squamous Cell Carcinoma (HNSCC) cell lines in control (0 Gy) and irradiated (48 h after 10 Gy) condition (Fig. 2A, S4).

**Fig. 2.**
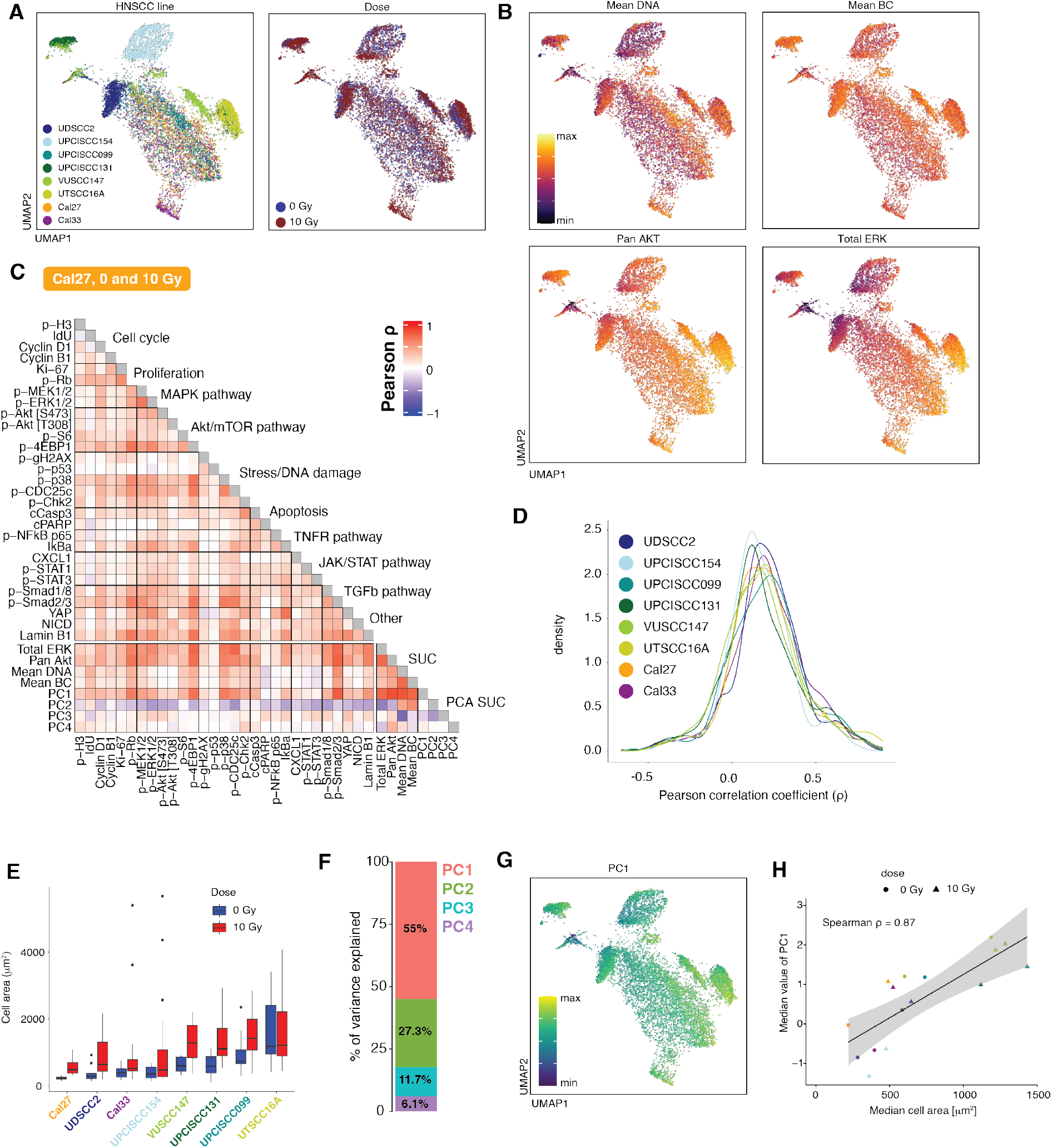
Unwanted covariance in a mass cytometry data set coming from heterogeneous cell area and other factors. **A)**. UMAP coloured by cell line (left) and dose (right) calculated by excluding cell cycle and proliferation markers (n = 800 cells per line and dose). **B)**. UMAP is coloured by the expression values of the 4 SUCs: mean DNA (top left), mean BC (top right), pan Akt (bottom left), and total ERK (bottom right). Expression values were asinh-transformed and min-max normalised. **C)**. Pearson correlation coefficients between asinh-transformed and z-scored expression values of markers in the Cal27 cell line across cells from 0 and 10 Gy. **D)**. Distribution of Pearson correlation coefficients between markers in all cell lines across cells from 0 and 10 Gy. **E)** Measurements of the cell area via microscopy images before fixation (n = 16.5 *±* 3.7 cells per cell line and condition, Fig. S4B). **F)**. Percentage of variance explained by each PC of a PCA based on the SUCs. PCA was calculated based on the asinh-transformed and z-scored expression values of the 4 SUCs in B). **G)**. UMAP coloured by PC1 of a PCA based on the 4 SUCs. PC1 values were min-max normalised. **H)** Spearman correlation coefficients between the median cell area (*µm*^2^) from E) and median value of PC1 from G) per cell line and dose.

Since cell volume and labelling efficiency cannot be directly measured with mass cytometry, we use four <u>S</u>urrogates of <u>U</u>nwanted <u>C</u>ovariance (SUCs, Eq. S1): (1) Mean DNA: Mean value of normalised iridium channels (Eq. S2), (2) Mean BC: Mean value of the highest (used) barcoding isotopes per cell (Eq. S3), (3) pan Akt, and (4) total ERK (Fig. 2B).

Iridium is a common DNA stain in mass cytometry. We noted that Ruthenium, previously proposed by Rapsomaniki et al. [2018] to correct for the cell volume, strongly correlates with DNA staining (Fig. S3A).

Mass cytometry is often performed with multiplexed samples which are stained with a specific combination of isotopes, *e*.*g*.: palladiums or telluriums [Zunder et al., 2015; Willis et al., 2018], acting as a barcode. These barcoding reagents bind unspecifically to surface proteins and, when stained while cells are fixed, also intracellular proteins [Zunder et al., 2015]. Thus barcode signals may be used as a surrogate of cell volume.

Total ERK and pan Akt are commonly used as loading controls for normalization in *e*.*g*.: Western blotting experiments, as they are typically abundant proteins that are relatively stable under different experimental conditions.

These four SUCs strongly correlate with each other and to the majority of the markers in all studied HNSCC cell lines (Fig. S5, S6). The correlations for the Cal27 cell line are depicted in Fig. 2C as an example. Some correlations are authentic and expected, such as between proliferation markers (p-Rb and Ki-67), members of the MAPK pathway (p-MEK1/2 and p-ERK1/2), regulators of cell cycle progression and protein synthesis (p-Rb and p-4EBP1), proteins in the DNA damage response pathway (p-p53 and p-γH2AX), and proteins involved apoptotic cell death (p-Chk2, cCasp3, cPARP, NF-κB and IκBα). However, other correlations are suspicious and most likely driven by unwanted covariance, especially between p-Rb, p-38, p-CDC25c, p-Smads, YAP and the majority of the measured markers (Fig. 2C). This distribution of correlation coefficients is observed in all the studied HNSCC lines (Fig. 2D, S5, S6)

We quantified the cell area via microscopy images for each HNSCC cell line and condition (Fig. 2E, Fig. S4A, B). We observed variations in the median cell area, both across different cell lines within the same condition and for each cell line across radiation conditions. The median area of the cells increased after irradiation with 10 Gy, which agrees with previously observed cellular enlargement of irradiated cells [Rene and Nardone, 1968; Ronny Sham et al., 2020; Ren et al., 2023]. The UPCISCC131 cell line displayed the highest increase in median cell area following irradiation.

PCA calculated on the four SUCs shows that more than half of the variance across SUCs is explained by PC1 (Fig. 2F,G), which strongly correlated with the quantified median cell area (*ρ* = 0.87, 2H), in contrast with subsequent PCs (Fig. S6D).

In an uncorrected mass cytometry data set, these differences in cell area (and therefore marker abundance) and other factors like staining efficiency will confound the comparisons between cell lines and perturbations, masking meaningful biological information.

### 2. RUCova enables improved classification of apoptotic cells by uncovering previously obscured cell populations

To remove the unwanted covariance coming from heterogeneous cell area, but also potentially from heterogeneous staining efficiency, we applied the RUCova method using the interaction model 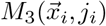 with the cell lines as samples (*j*).

After removing the covariance with all SUCs (*i*.*e*, using all four PCs as predictive variables) spurious correlations were removed and authentic correlations were kept (Fig. 3A-C), enabling differentiation between activated (apoptotic and MAPK pathway), and non-activated signalling pathways (JAK/STAT and TGFβ). The covariance between marker abundances was substantially decreased in all HNSCC cell lines, especially after regressing out the correlation with PC1 and PC2 (Fig. 3D, S5, S6). Consistently, we observed a general decrease in the standard deviation (*σ*) of the signals for each condition after applying RUCova (Fig. 3E). However, for some markers, we observed an unexpected increase in the standard deviation relative to the original distributions (arrows in Fig. 3E). For IdU in the UPCISCC154 line, p-γH2AX in the UPCISCC131 line, and p-p53 in the Cal33 line, the higher standard deviation observed after RUCova was attributed to the assignment of non-zero values to the artificial zero values typically present in a mass cytometry measurement (Fig. 3F). The rise in the standard deviation of the distribution of the apoptotic marker cleaved Caspase-3 (cCasp3) in the irradiated UPCISCC131 line can be attributed to two factors: the assignment of non-zero values and an increased dissimilarity in cCasp3 signal between the non-apoptotic and apoptotic populations (Fig. 3F,G). We categorized the apoptotic cells in the irradiated UPCISCC131 cell line by analyzing their cCasp3 signals and establishing decision thresholds before and after RUCova (based on PC1 to PC4) (dashed vertical lines in Fig. 3G). The classification based on the data after applying RUCova allowed us to increase the identification capabilities of apoptotic cells by 47.5 % (Fig. 3H). Before applying RUCova, Apoptotic cells showed lower original cCasp3 signals, making them less distinguishable from non-apoptotic cells. This originally lower cCasp3 signal corresponded to lower PC1 values (Fig. 3I).

**Fig. 3.**
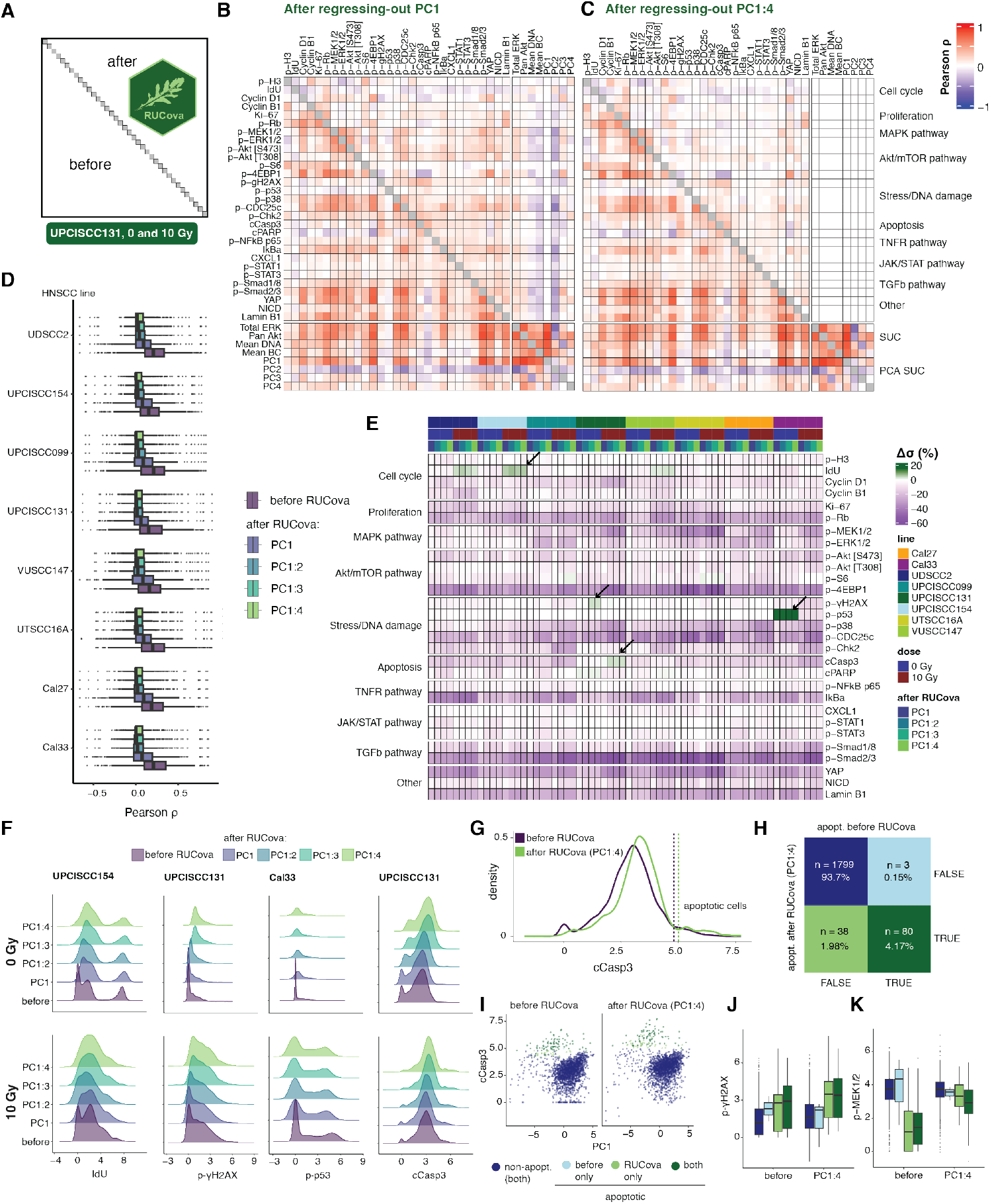
RUCova removes unwanted covariance and allows improved classification of apoptotic cells. **A)** Scheme of the correlation heatmap, where the lower and upper triangles show the Pearson correlation coefficients between marker values before and after RUCova (using the interaction model 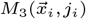 per cell line), respectively. Diagonal unity values are depicted in grey. Pearson correlation coefficients were calculated between the asinh-transformed and z-scored expression values of markers. **B**,**C)** Correlation heatmap with the upper triangle showing the Pearson correlation coefficients between marker values in the UPCISCC131 cell line across cells from 0 and 10 Gy after applying RUCova based on **B)** PC1, **C)** PC1 to PC4. **D)** Boxplots for Pearson correlation coefficients between markers per cell line across cells from 0 and 10 Gy, based on the expression values before and after RUCova. **E)** Heatmap of percentual difference in the standard deviation *σ* of each marker’s distribution after applying RUCova relative to the distributions before RUCova 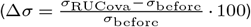 per cell line and dose levels. Arrows indicate specific cases where Δ*σ >* 0. **F)** Density plots for asinh-transformed expression values of IdU, p-*γH*2*AX*, p-p53, and cleaved Casp3 before and after RUCova (y-axis) in cell lines where Δ*σ >* 0 (arrows in panel E) for 0 Gy (first row) and 10 Gy (second row). **G)** Density plot for asinh-transformed expression of cleaved Casp3 in the irradiated (10 Gy) UPCISCC131 cell line, before (purple) and after RUCova based on all four PCs (green). Dashed vertical lines indicate the decision thresholds for apoptotic (higher values of cleaved Casp3) and non-apoptotic populations according to the cleaved Casp3 distribution before and after applying RUCova. **H)** Confusion matrix for classification of apoptotic cells in the irradiated UPCISCC131 cell line before and after RUCova (based on all four PCs) according to the decision thresholds in G). **I)** Scatter plots of PC1 vs. asinh-transformed values of cleaved Caspase-3 before (left) and after RUCova (right) for irradiated cells in the UPCISCC131 line. Cells are coloured by apoptotic status. **J**,**K)** Boxplots of asinh-transformed expression of **J)** p-γH2AX and **K)** p-MEK1/2 before and after RUCova in irradiated UPCISCC131 cells according to their apoptotic status.

After RUCova, the expected differences in markers like the DNA-damage marker γH2AX between apoptotic and nonapoptotic cells became discernible (Fig. 3J, S7), while potential artefacts, like lower p-MEK1/2 signals in apoptotic cells, were reduced (Fig. 3K, S7).

The use of RUCova in this data set enabled the preservation of authentic correlations while removing spurious ones, facilitating the differentiation between activated and non-activated signalling pathways after irradiation in different HNSCC lines. The application of RUCova allowed a clearer understanding of apoptotic marker distribution and its relation to cell size reduction, highlighting its efficacy in elucidating complex data patterns in mass cytometry analysis.

### 3. RUCova enhances the reliability of perturbation comparisons by eliminating cell size artefacts

To understand how cell size and other factors can confound mass cytometry data and especially analyses of cellular responses, we perturbed cells from the HNSCC Cal33 cell line using EGF stimulation (30 min), EGFR inhibition (Gefitinib, 24 h), Etoposide treatment (2 h), IFN-β stimulation (30 min), IGF stimulation (30 min), PI3K inhibition (24 h), and starvation alone (24 h). We then sorted the cells into two groups based on size (small and large cells) using fluorescence-activated cell sorting (FACS) (Fig. 4A).

**Fig. 4.**
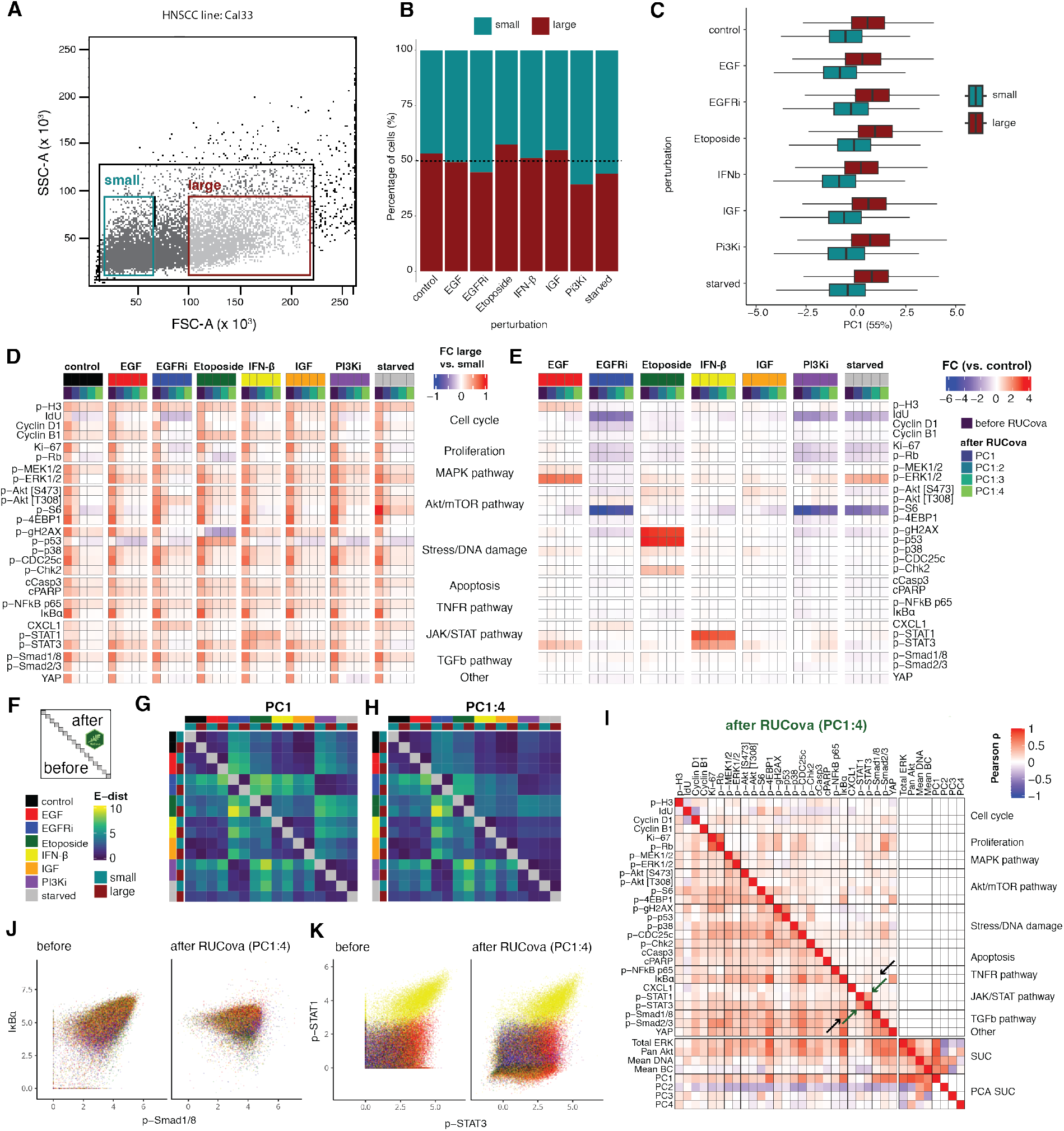
RUCova successfully removes cell size artefacts enabling accurate perturbation analyses. **A)** Scatter plot of the FACS parameters SSC-A vs FSC-A. Gates for size-sorted groups of small (blue) and large (red) cells in the Cal33 cell line before mass cytometry measurement. **B)** Percentage of small and large cells per perturbation condition in the Cal33 cell line. **C)** Boxplots of PC1 (of a PCA based on the four SUCs) per perturbation and size group. **D)** Fold Change (FC) of asinh-transformed values between large and small cells in all conditions, before and after applying the simple RUCova model 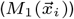 using different numbers of PCs. **E)** Fold Change (FC) of asinh-transformed values between perturbation and control condition, before and after applying the simple RUCova model 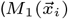 using different number of PCs. **F)** Scheme of the E-distance heatmap, where the lower and upper triangles show the E-distance between conditions before and after RUCova (using the simple model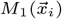, respectively. **G**,**H)** E-distance heatmap between 1000 cells per condition and sorted population for data after applying RUCova using **G)** only PC1, **H)** PC1 to PC4. **I)** Correlation heatmap with the upper triangle showing the Pearson correlation coefficients between marker values across all perturbations and sorted populations after RUCova based on PC1 to PC4. Black arrows indicate an artifactual correlation which is removed after applying RUCova, and green arrows indicate a real correlation which is kept. **J, K)** Scatter plots of Cal33 cells coloured by perturbation before (left) and after (right) applying RUCova based on all four PCs. **J)** asinh-transformed values of p-Smad1/8 vs. IκBα (artifactual correlation). **K)** asinh-transformed values of p-Stat1 and p-Stat3 (real correlation driven by IFN-β-stimulated cells).

Overall, most of the perturbations resulted in similar proportions of sorted cells, as shown in Fig. 4B. However, in the case of PI3K inhibition, most sorted cells were smaller. The PC1 of a PCA based on the SUCs was substantially higher for large compared to small cells (Fig. 4C). Correspondingly, average marker values were consistently higher in large cells compared to small cells across all perturbations (Fig. 4D), illustrating once again how cell size confounds mass cytometry data. Upon applying RUCova using the simple model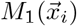, these differences in mean values were notably reduced, particularly after removing correlations with all four PCs. For certain markers and perturbations (e.g., p-p53 in Etoposide treatment and p-STAT1 in IFN-β stimulation) considerable fold changes between large and small cells persisted after RUCova, indicating the method’s ability to preserve genuine biological signals while eliminating artificial ones. Across large and small cells, the fold changes between perturbation and control conditions were usually maintained after applying RUCova (Fig. 4E). Nevertheless, the removal of unwanted covariances led to modified fold-changes in some cases: p-ERK1/2 was increased after PI3K inhibition relative to control, which might indicate a compensatory signalling mechanism as a means of maintaining cell survival and proliferation. These reductions or increments in the fold-change after RUCova could also be due to the unbalanced proportion of small and large cells in a perturbation compared to the control condition. By assessing the pairwise E-distance (Peidli et al. [2024]) between perturbation conditions and sorted populations both before and after applying RUCova, we observed that differences between small and large cells were effectively eliminated, along with dissimilarities arising from variations in cell size distribution, such as those seen in the PI3K condition compared to the others. We also observed small increments in the E-distance after implementing RUCova, especially between EGFR inhibition and starvation (Fig. 4F,G,H, Fig. S8A).

The Pearson correlation coefficients between markers across the entire data set after RUCova provide reliable information. An example of a regressed-out artefactual correlation is p-Smad1/8 and IκBα (black arrow Fig. 4I,J), which was mainly driven by the size of the cells (Fig. S8C). An example of a real correlation that was kept after RUCova is between p-Stat1 and p-Stat2 mainly driven by IFN-β stimulation (green arrow Fig. 4I,K, S8D).

This suggests that RUCova successfully reduced the dissimilarities between conditions due to cell size and other potential sources of unwanted covariance, providing a clearer understanding of the underlying biological responses to different stimuli.

## Discussion

Mass cytometry data is contaminated by variance that is induced by variations in cell size, staining efficiency and due to other technical artefacts, leading to spurious correlations between markers. Here we describe RUCova, a method to regress out such unwanted co-variation. The method consists of fitting a model for each marker based on Surrogates of Unwanted Covariance (SUCs), such as mean DNA, mean barcode signal and total protein markers such as total ERK and AKT. Previous approaches used fixed relation between marker abundance and cell size stains [Rapsomaniki et al., 2018] yielding sub-optimal results, as the extend of correlation between protein abundance and cell size varies and depends *e*.*g*. on protein localisation [Lanz et al., 2023].

Cell size can exhibit variability across distinct cell lines, cell types (e.g., various immune cells), and even within different tissue microenvironments [Liu et al., 2022]. If the association between cell size and protein abundance varies, RUCova can incorporate this using the interaction model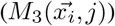, where the slopes of the regression fit can account for differences among samples (cell lines, cell types, or cell clusters).

In Fig. 2, S5, S6 we illustrated that the Pearson correlation coefficients between markers display consistently high values, posing challenges in distinguishing between activated and non-activated signalling pathways. The majority of these markers exhibited strong correlations with the SUCs, especially with the first principal component (PC1) across SUCs which explained 55 % of the variance and highly correlated with cell area, thereby confirming the artifactual nature of these elevated correlations.

In Fig. 3 we demonstrated how spurious correlations and resulting variance in marker distributions were effectively mitigated with RUCova. We noticed that PC1 of the SUC represents cell area and that regressing out PC1 and PC2 removes spurious correlation effectively. We recommend using PCA components on the SUC as predictors in the model. By adjusting the number of PCs used, the user of RUCova can adjust the extent of unwanted correlation to be removed. PC1 may also serve as a proxy of cell size in later analysis. Mass cytometry data often contains an inflation of zero values that are technically induced by the sensitivity thresholds of the instruments. While RUCova widely reduces variance within conditions, we observed that the variance of markers with a large number of zeros increased as those will be non-zero values after regression. Some approaches developed for mass cytometry data address these zero values by imputing them with estimated values [Li et al., 2017; Minoura et al., 2020] or by extrapolating measurements from other panels using k-nearest neighbour methods [Abdelaal et al., 2019]. However, applying these imputation methods to uncorrected data may introduce bias, as the resulting non-zero values could be influenced by existing covariance in the dataset. In contrast, RUCova addresses this issue by assigning non-zero values while simultaneously removing such covariance.

An important application of RUCova is to identify populations of cells that may be obscured due to heterogeneous cell sizes. In this article, we showed that RUCova allowed us to identify about 50% more apoptotic cells in a data set. These previously hidden cells exhibited the lowest PC1 values, suggesting they were smaller in size, a known phenomenon in apoptotic cells which undergo cell shrinkage during the early stages of apoptosis [Kerr et al., 1972; Jänicke et al., 1998; Albeck et al., 2008]. Importantly, RUCova also corrected marker expression for apoptosis-related signals, including higher levels of the DNA damage marker p-*γH*2*AX* compared to non-apoptotic cells.

To directly compare how RUCova corrects the data between small and large cells, we employed fluorescence-activated cell sorting (FACS) to separate large and small cells. RUCova removed differences in signals between small and large cells with only very few biologically plausible exceptions. These include p-Stat1 which showed differences between large and small cells following IFN-β stimulation. IFN-β induces inflammation which leads to larger cell sizes [Han et al., 2022], thus higher p-Stat1 signal may be a genuine signal indicative of inflammation-induced cell enlargement.

RUCova relies on the selection of SUCs to model and remove unwanted covariances in the mass cytometry data. However, the effectiveness of RUCova may be influenced by the choice and availability of suitable SUCs. In some cases, identifying appropriate SUCs that adequately capture all sources of unwanted covariance may be challenging. RUCova operates under the assumption of linearity in the relationship between marker expression and SUCs. While this assumption may hold for many cases, there may be instances where the relationship between markers and SUCs is non-linear, leading to potential limitations in the accuracy of the correction performed by RUCova. The performance of RUCova may be sensitive to the specific modelling approach chosen (simple, offset or interaction model), the selection of the number of principal components or the decision to use uni– or multi-variate modelling. Suboptimal model selection may lead to incomplete correction of unwanted covariances or overfitting of the data, potentially introducing biases or artefacts into the analysis. We recommend employing and comparing multiple modelling approaches. While RUCova aims to improve the interpretability and reliability of mass cytometry data by removing unwanted covariances, it is essential to consider the potential impact of data correction on biological interpretation. Over-correction or removal of genuine biological signals alongside technical artefacts may obscure meaningful biological insights or introduce biases into downstream analyses. For optimal utilization of RUCova and accurate interpretation of its results, it’s recommended that mass cytometry users visually inspect cell morphology and estimate cell size. Additionally, verifying any correlation between cell size estimates and covariance levels in the dataset is essential. This approach ensures a comprehensive understanding of the data and boosts confidence in the regression process by identifying potential sources of unwanted variability.

## Conclusion

In conclusion, our study introduces RUCova as a powerful tool for removing unwanted covariance in mass cytometry data, thereby enhancing the accuracy and reliability of downstream analyses. By effectively addressing technical artefacts associated with heterogeneous cell size and staining efficiency, RUCova facilitates the uncovering of genuine biological signals and contributes to a deeper understanding of cellular processes. Our findings demonstrate the utility of RUCova in elucidating complex data patterns, facilitating the identification of activated signalling pathways, and improving the classification of apoptotic cells. Furthermore, we emphasize the importance of thoughtful model selection and validation, as well as the critical interpretation of results in the context of biological insights. Moving forward, continued refinement and validation of RUCova and related methodologies will further enhance their utility in advancing our understanding of cellular biology and disease mechanisms.

## Supplementary data

Supplementary material is available online (PDF).

## Availability and implementation

R package is available on https://github.com/molsysbio/RUCova. Detailed documentation, data, and the code required to reproduce the results are available on https://doi.org/10.5281/zenodo.10913464.

## Competing interests

None declared.

## Funding

The funding for this work was provided by the BMBF grants number 02NUK047C & 02NUK047E (ZiSS-Trans), 02NUK086A & 02NUK086E (Senirad), 16LW0239K (MSTARS-2), and DFG research training group RTG2424 (CompCancer).

## Author contributions statement

R.A.G. and N.B. conceptualized the project. T.S., S.M., and A.S. conducted the experiments. R.A.G developed the RUCova method and R package in collaboration with T.S. and N.B. R.A.G. and N.B. wrote and reviewed the manuscript. N.B. and K.L. provided supervision.

## Acknowledgments

We acknowledge the BIH Cytometry Core Facility for their help with cell sorting and mass cytometry data acquisition.

## Notes

https://github.com/molsysbio/RUCova

### Competing Interest Statement

The authors have declared no competing interest.

### Summary of Updates

We corrected spelling and grammatical errors. Additionally, the background of the PNG figures was changed from transparent to white.

