## Supplementary material for "RUCova: Removal of Unwanted Covariance in mass cytometry data"

### Supplementary Material for
### RUCova: Removal of Unwanted Covariance in mass cytometry data

Rosario Astaburuaga-García, Thomas Selll, Samet Mutlu, Anja Sieber, Kirsten Lauber, Nils Blüthgen

R package is available on https://github.com/molsysbio/RUCova. Detailed documentation, data, and the code required to reproduce the results are available on https://doi.org/10.5281/zenodo.10913464.

**S3. HNSCC data set** **9**
**S4. FACS-sorted data set** **11**

### S1. The RUCova method

RUCova comprises two major steps. First, it fits a multivariate model for each measured marker ($m$) across cells ($i$) from samples ($j_i$) with respect to the surrogates of unwanted covariance (SUC) $\vec{x_i}$ (Eq. **??**). Second, it eliminates such dependency by assigning the residuals $\epsilon$ of the model as the new modified expression of the marker (Eq. **??**).

#### S1.1 Surrogates of Unwanted Covariance ($\vec{x_i}$)

Since cell volume and labeling efficiency cannot be directly measured with mass cytometry, we use four Surrogates of Unwanted Covariance (SUCs): (1) Mean DNA: Mean value of normalised iridium channels, (2) Mean BC: Mean value of the highest (used) barcoding isotopes per cell, (3) pan Akt, and (4) total ERK.

$$\vec{x_i} = [\mathrm{mean\ DNA_i}, \mathrm{mean\ highest\ BC_i}, \mathrm{total\ ERK_i}, \mathrm{pan\ Akt_i}] \tag{S1}$$

Mean DNA per cell $i$ was calculated as:

$$\mathrm{mean\ DNA_i} = \frac{\mathrm{Ir191_{norm,i}} + \mathrm{Ir193_{norm,i}}}{2} \tag{S2}$$

where $\mathrm{Ir191_{norm}}$ and $\mathrm{Ir193_{norm}}$ are the percentile-normalised intensities (*e.g.,* to the 95th percentile) of the iridium intercalators serving as DNA stains for each cell $i$. The mean BC signal per cell $i$ was calculated as:

$$\mathrm{mean\ highest\ BC_i} = \frac{1}{\mathrm{N_{BC}}} \sum_{\mathrm{k=1}}^{\mathrm{N_{BC}}} \mathrm{BC_{i,k,norm}} \tag{S3}$$

where $\mathrm{BC_{norm}}$ are the percentile-normalised intensities of the barcoding isotopes across all cells (*e.g.,* to the 95th percentile), $\mathrm{N_{BC}}$ is the number of barcoding isotopes used per cell (e.g.: $\mathrm{N_{BC}} = 3$ for the Fluidigm kit of Palladium isotopes), and $\mathrm{BC_k}$ is the barcoding isotope with the k-th highest signal in cell $i$, meaning the isotope was used in that cell to barcode it. The surrogates pan Akt and total ERK are markers included in our mass cytometry panel.

By taking the zero-centered distributions of the SUCs ($x_i^c = x_i - \frac{1}{N_i}\sum_i^{N_i} x_i$), the mean values of the markers $m$ across all cells are kept after applying RUCova (Fig. **??**). If a more conservative approach is desired where the fold changes between samples should be kept, each SUC should be centered per sample. This approach is illustrated in Fig. S1) for the three RUCova models : simple (Eq. **??**), offset (Eq. **??**), and interaction (Eq. **??**). Similarly, when using PCs as the predictive variables, SUCs can be z-score normalised by sample before performing PCA.

Performing PCA on the SUCs may be useful, especially to better identify axes correlating with confounding factors such as PC1 and cell size (Fig **??**H).

#### S1.2 RUCova parameters

Here we illustrate the application of RUCova using the Head-and-neck squamous cell carcinoma (HNSCC) data set (from Fig. **??**, **??**), for which we choose the interaction model $M_3(\vec{x_i}, j_i)$ (Eq. **??**). Fig. S2A shows the distribution of the SUCs. PC1 (of a PCA on the SUCs) was not only dependant on one SUC but rather on the combination (loadings in Fig. S2B).

One of the outputs of RUCova is the adjusted R-squared ($\mathrm{R^2_{adj}}$) or coefficient of determination, which is a useful metric to evaluate the goodness of fit (Fig. S2C). It quantify the proportion of the variance in the dependent variable (marker) that is explained by the independent variables (surrogates) in the model. The Adjusted R-squared is *adjusted* by the number of independent variables used for predicting the target variable. This is done to account for the automatically increase of $R^2$ values when extra explanatory variables are added to the model. By analysing the $\mathrm{R^2_{adj}}$,we can determine whether adding new variables to the model actually increases the model fit.

$$R^2_{adj} = \Big(1 - \underbrace{(1 - \frac{SS_{res}}{SS_{tot}})}_{R^2}\Big) \cdot \frac{n-1}{n-q-1} \tag{S4}$$

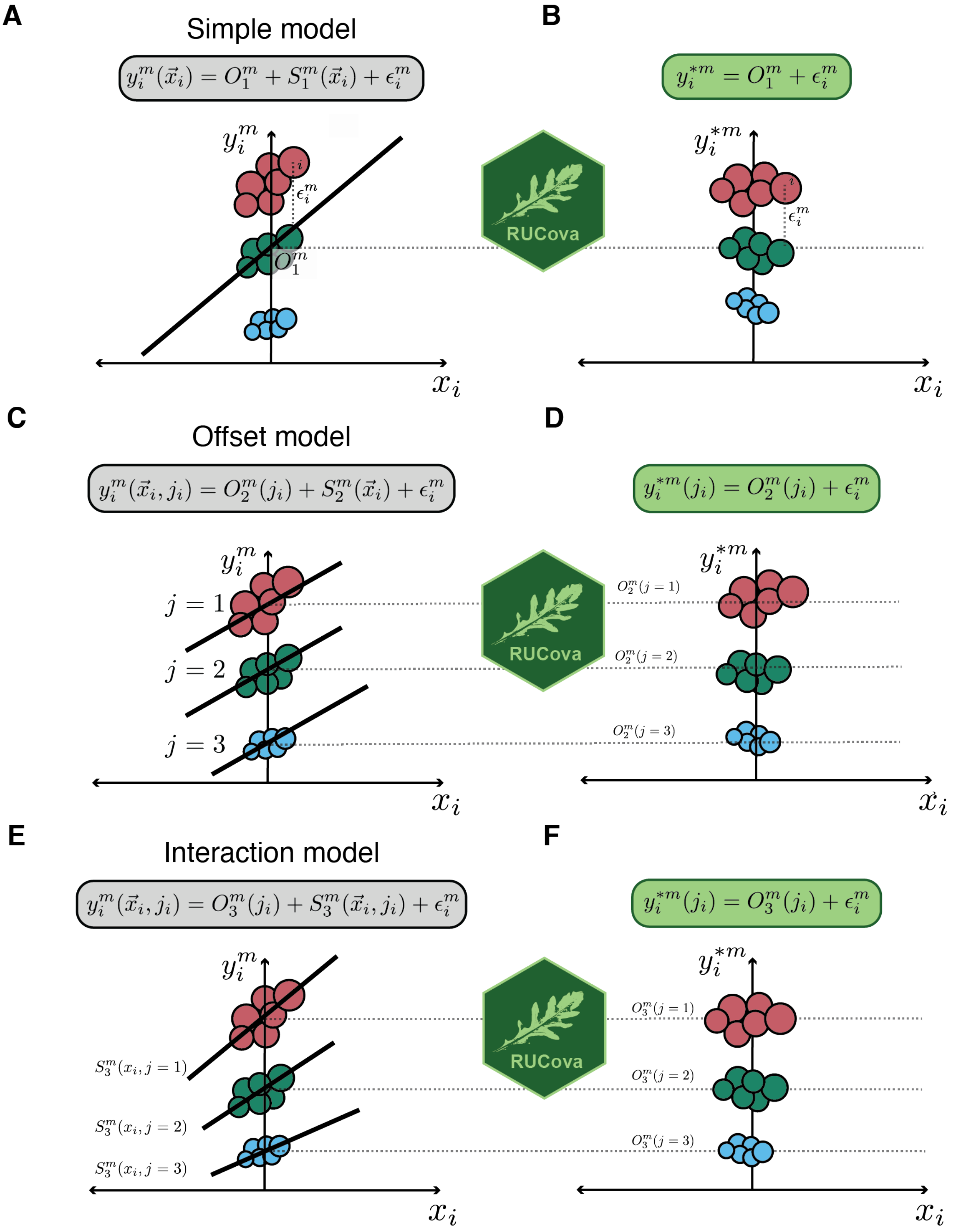

Figure S1: **Illustration of the RUCova method and its three different models, when surrogates are centered per sample. A, C, E)** Original expression $y_i^m$ of a marker $m$ before RUCova as a function of a centered expression of a SUC or PC. Illustrative regression line and equation corresponding to each model. **B, D, F)** Modified expression $y_i^{*m}$ of a marker $m$ after RUCova. **D, F)** Keeping the offset $O^m(j_i)$ between samples $j$. **A, B)** Simple model: one fit across the input data set with intercept $O_1^m$ and residuals $\epsilon_i$. **C, D)** Offset model: one slope $S_2^m(\vec{x}_i)$ for the whole input data set and different intercepts $O_2^m(j_i)$ between samples. **E, F)** Interaction model: one fit per sample $j$ with intercepts $O_3^m(j_i)$ and slope $S_3^m(\vec{x}_i, j_i)$.

where $SS_{res} = \sum_i \epsilon_i^2$ is the residual sum of squares, $\epsilon_i$ are the residuals of the model, $SS_{tot} = \sum_i (y_i - \overline{y})$ is the total sum of squares, $\overline{y}$ is the mean value of the marker, $n$ is the sample size (total number of cells $i$) and $q$ is the number of explanatory variables in the model.

The model coefficients are also included in the output of RUCova. Assessing the slope coefficients is useful to better understand the effect size of each SUC on the markers. We standardised the slope coefficients in order to make them comparable. In the case of the interaction model, where the slopes $\alpha^m$ depend on samples $j_i$ and the SUC $p$, we standardised the slope by multiplying it by the standard deviation of the corresponding SUC in each sample $j$

$(\sigma_{x_{j_i,p}})$ and dividing it by the standard deviation of the marker $m$ in sample $j$ $(\sigma_{y^m_{j_i}})$ :

$$\alpha^{m*}_{j_i,x_{i,p}} = \alpha^{m}_{j_i,x_{i,p}} \cdot \frac{\sigma_{x_{j_i,p}}}{\sigma_{y^m_{j_i}}} \tag{S5}$$

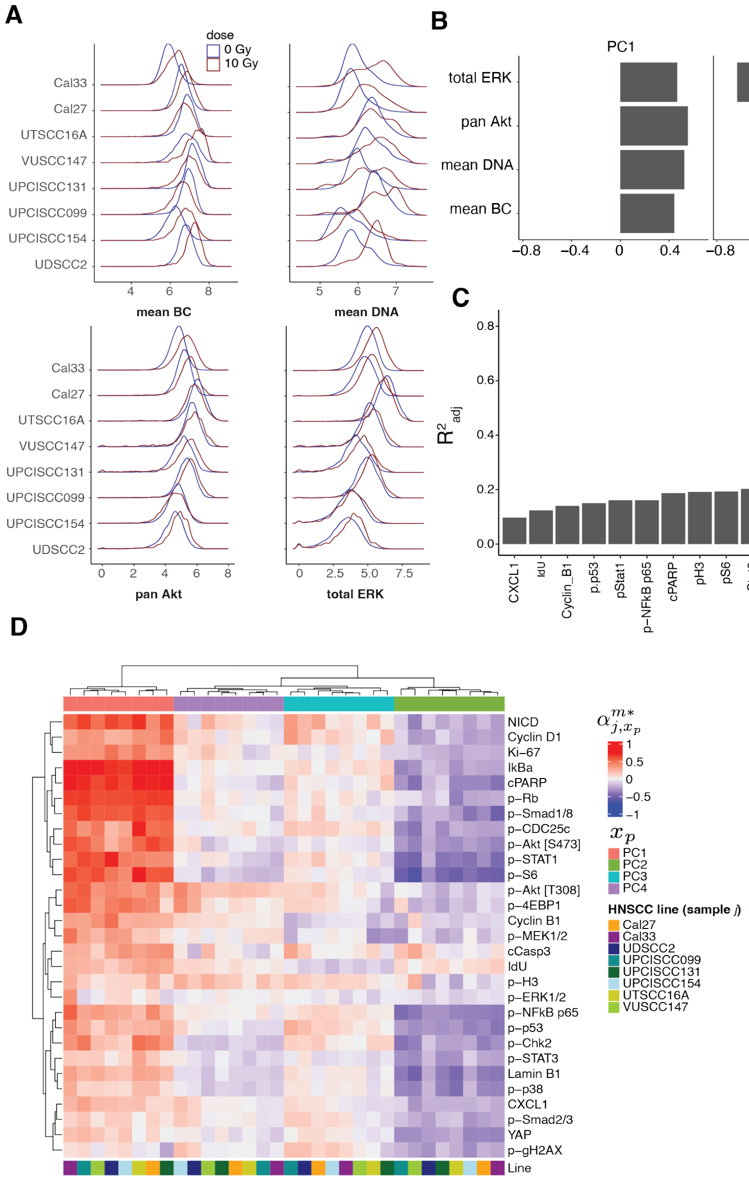

Figure S2: **A)**. Asinh-transformed values of surrogates of unwanted covariance (SUC) across cell lines and irradiation conditions. **B)**. Loadings of a PCA based on asinh-transformed and z-score normalised SUCs. **C)**. Adjusted R-squared for each marker after applying the interaction model. **D)**. Standardised slope coefficients for each marker after applying the interaction model with cell lines as samples.

#### S1.3 Alternative SUCs

Other markers may be used as SUCs, for example, the Ruthenium isotopes proposed by Rapsomaniki et al. [2018]). However, they might not add additional information as we observed a high correlation with the mean DNA values (Fig. S3A) in their published data set. They proposed normalising the marker intensity by the mean Ruthenium staining per cell. Consistently, dividing by the mean DNA leads to very similar normalised values (Fig. S3B, first and second heatmap). This approach assumes the same relationship between all markers and cell volume, which might not be correct. Consequently, many correlations between markers and ruthenium or mean DNA are kept after normalisation. On the contrary, by regressing-out the correlations with Ruthenium or DNA (Fig. S3B, third to fifth heatmap) with RUCova and allowing different relationships between cell volume and marker abundance for different markers, all the correlations with ruthenium and DNA stainings were removed. We observed similar results with our experimental measurements of Ruthenium staining in Cal33 cells (Fig. S3C). By including pan Akt and total ERK as SUCs we were able to remove unwanted covariance at a greater extent, especially in the signals of p-Smad2/3 and YAP.

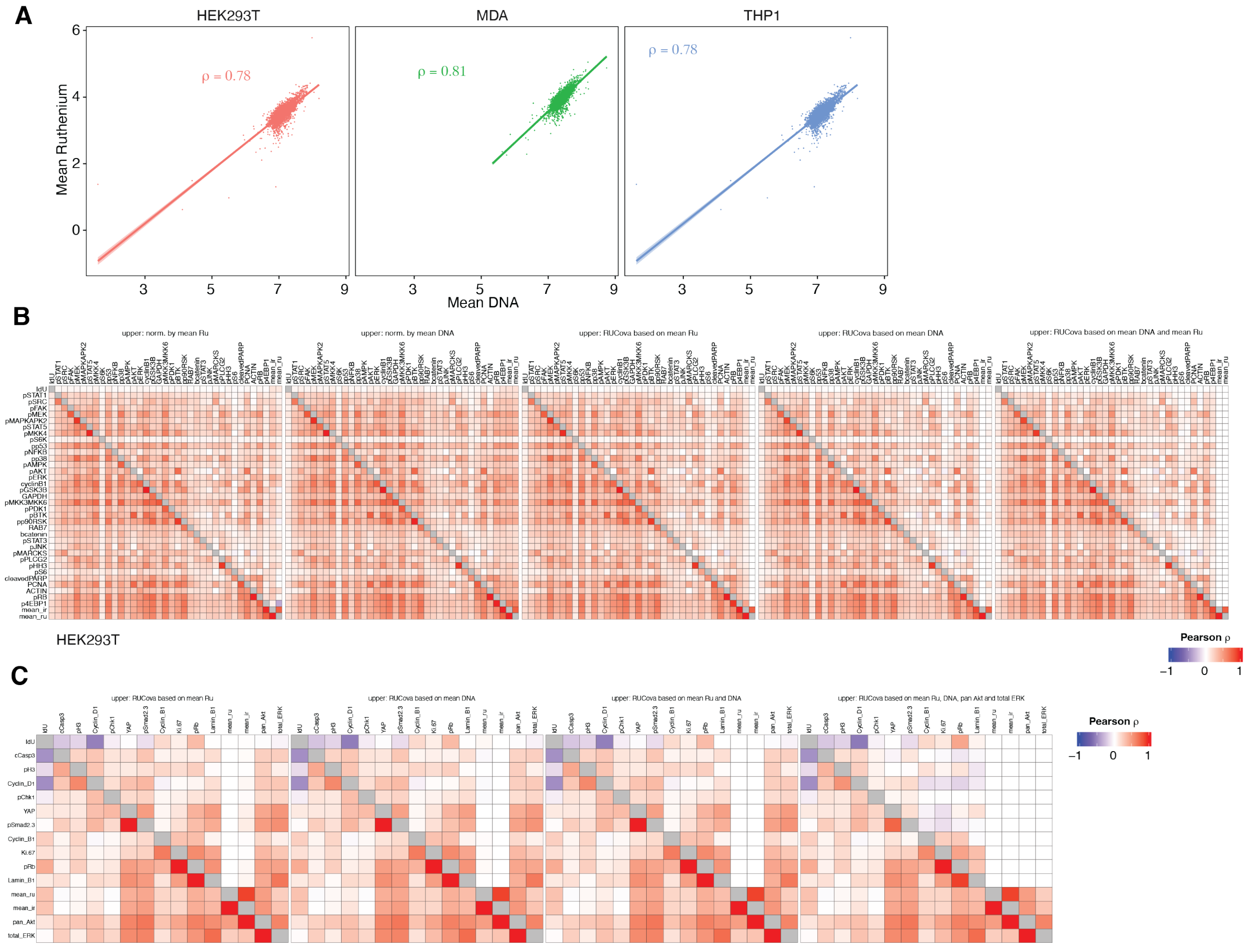

Figure S3: **A)** Scatter plots of asinh-transformed signals of mean Ruthenium and mean DNA staining in three different cell lines (Data from [Rapsomaniki et al., 2018]). **B)** Pearson correlation coefficients between marker abundances in the HEK293T cell line (Data from [Rapsomaniki et al., 2018]) before any normalisation (lower triangle) and after the normalisation or regression indicated on the top of each heatmap. **B)** Pearson correlation coefficients between marker abundances in the Cal33 cell line before RUCova (lower triangle) and after RUCova (upper triangle) following based on the SUCs indicated on the top of each heatmap.

### S2. Experimental details

#### S2.1 Cell culture

Cells from 10 HNSCC cell lines (UDSCC2, UPCISCC040, Cal27, UPCISCC099, UPCISCC131, UPCISCC154, UMSCC1, VUSCC147, UTSCC16A, Cal33) were cultured in DMEM supplemented with 10 % FCS, 1 % Glutamax (Gibco, 35050061), and 1 % Penicillin/Streptomycin (Gibco, 15140122). Cells were seeded into 6-well plates at day -1 (for 0 Gy 400.000 cells/well, for 10 Gy 700.000 cells/ well). Right before radiation treatment at day 0, medium was replaced with fresh medium, and cells were irradiated with 10 Gy using an RS225 X-ray cabinet (X-Strahl, Camberley, UK) operated at 200 kV/10 mA (Thoraeus filter, 1 Gy in 63 s), or left untreated. At 48 hours, the cells were prepared for fixation as follows: 30 min prior to fixation, IdU was added to the cell culture medium for a final concentration of 10 µM (1:5000 from stock solution; Standard Biotools, 201127), plates were rocked well but gently, and incubated at 37 °C for 30 min. Culture medium was discarded and plates washed once with PBS. Cell-ID™ Cisplatin (2:1000 from stock, 2 µM in PBS; Standard Biotools, 201064) was added to cells for 5 min at 37 °C and cells were washed once with culture medium (full medium including FCS) followed by 1x PBS washing. Fixation of cells was performed using methanol-free formaldehyde (2 % in PBS; stock from Pierce™ 16 %, 28906) at 37 °C for 15 minutes. Reaction was stopped by adding protein-containing medium followed by 2x PBS washing. Subsequently, cells were dissociated by adding Accutase for 45 min at 37 °C, scraping, pipetting, and straining through Flowmi (40 µm, BAH 136800040, Sigma) filter. Cells were transferred into low-binding reaction tube. After spinning cells down at 800 g for 5 min and discarding the supernatant, cells were re-suspended in 500 µl PBS/BSA (10 %) + 10 % DMSO and stored at −20 °C.

For cell size determination, same procedure was followed (without IdU/Cisplatin administration) including fixation and washing steps. Then, images were acquired through 6-well plate using Zeiss AxioObserver Z1 inverted microscope (Fig. S4A). The acquired images were then analyzed for cell area determination using ZEN 2.3 software and embedded toolkit. A minimum of 12 cells were marked for each condition (Fig. S4B). After pre-processing of mass cytometry data, two cell lines (UMSCC1 and UPCISCC040) were excluded from the analysis due to low number of cells in the irradiated condition ($n < 500$, Fig. S4C).

For FACS sorting and ASCQ-Ru experiments, similar cell culture procedures were followed using only Cal33 cells, but plates were not irradiated. For the FACS sorting experiment, cells were perturbed in the following way: 24 h prior fixation, FCS was removed from the growth medium for all samples but one control. Perturbations were either Gefitinib for 24 h at 10 µM, EGF for 30 min at 25 ng/ml, IFN-β for 30 min at 25 ng/ml, Etoposide for 2 h at 42 µM, GDC0941 for 24 h at 1 µM, or IGF for 30 min at 100 ng/ml. IdU incubation was started in parallel as described above.

#### S2.2 Mass cytometry

Frozen samples were thawed at 37 °C and washed 1x in PBS. Free nucleic acids were digested by incubating each sample with Benzonase (Pierce, 88700; 100 U/ml in PBS). The Cell ID™ 20-Plex Pd Barcoding Kit (Standard Biotools, 201060) was used for sample barcoding according to manufacturer's instructions. Subsequent processing steps were performed on multiplexed cell pools. For experiments with more than 20 conditions to be barcoded, monoisotopic Cisplatin was used as an additional identifying factor of cells already pooled based on Pd barcodes. Multiplexed cells were permeabilised by chilling for 10 min followed by incubating with ice-cold methanol for 15 minutes. After 2x washes in Cell Staining Buffer (Standard Biotools, 201068), mass-tagged antibodies were added for 30 min at room temperature, followed by 2 more washes in Cell Staining Buffer. Iridium DNA intercalator (Standard Biotools, 201192A) was added at a concentration of 63 µM in PBS for 20 min at room temperature. Subsequently, the sample was washed once in PBS and stored overnight at 4 °C in methanol-free formaldehyde (2 % in PBS; stock from Pierce™ 16 %, 28906). On the following day some samples were treated with 25 µg/ml ASCQ-Ru in 0.1 M $NaHCO_3$. Other samples were divided into "small" and "large" cells via FACS sorting based on FSC-A/SSC-A ratios. All samples were washed 2x in doubly distilled water and filtered through a 10 µm cell strainer prior to acquisition in a CyTOF 2 (Helios upgrade) mass cytometer.

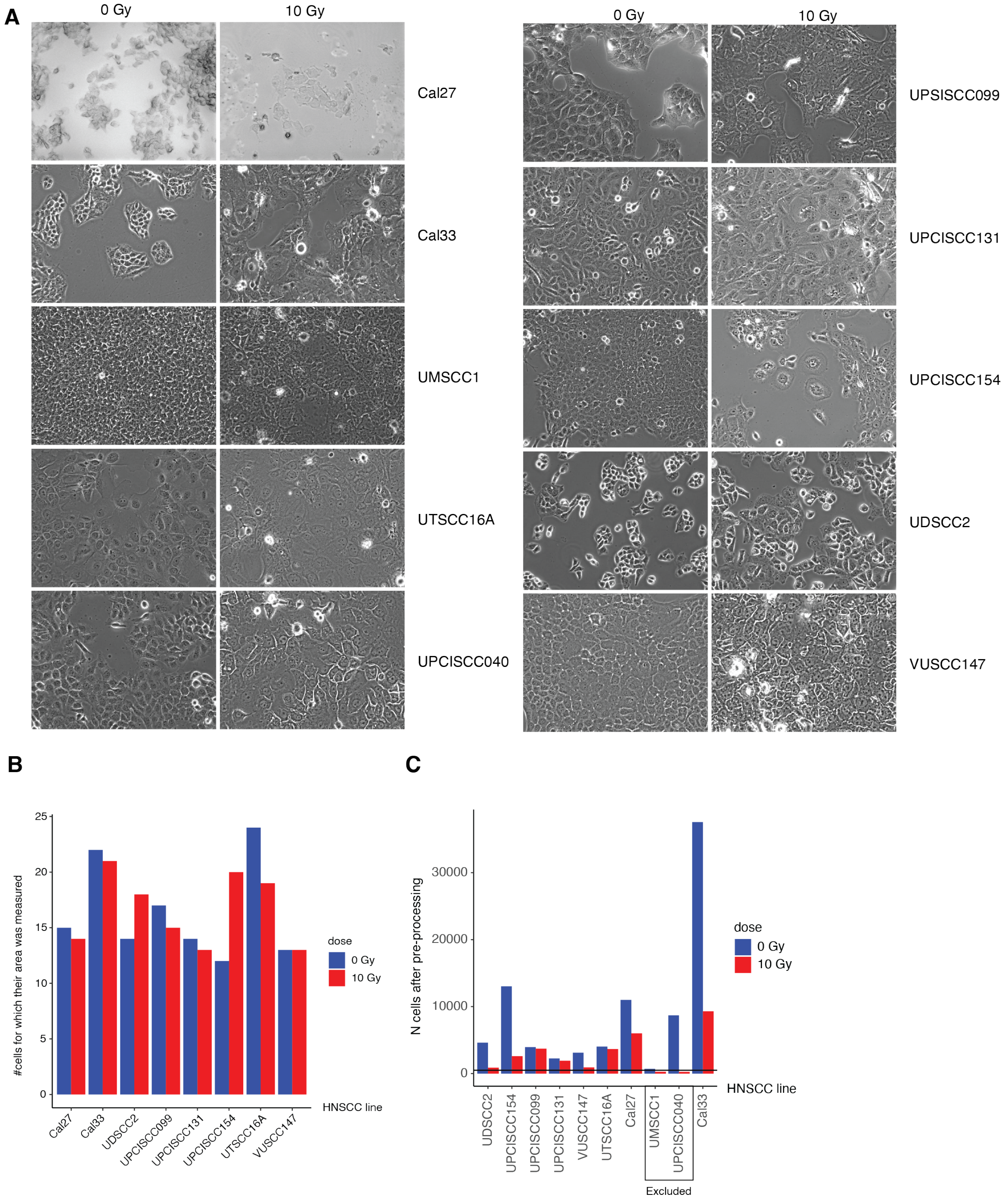

Figure S4: **A)** Microscopy images for the 10 HNSCC lines in each irradiated condition. **B)** Number of cells per cell line and condition for which the cell area was quantified. **C)** Number of cells per cell line and condition in the mass cytometry data set after pre-processing. UMSCC1 and UPCISCC040 were excluded due to low cell number ($n < 500$, horizontal line)

Table S1: Antibody panel for the three data sets: HNSCC, FACS-sorted and ASCQ-Ru

| label | Category | target | HGNC symbol | vendor | clone | HNSCC data set | FACS-sorted data set | ASCQ-Ru data set |
|---|---|---|---|---|---|---|---|---|
| 141Pr | Stress/DNA damage | pChk2 [T68] | CHEK2 | CST | C13C1 | X | X | |
| 142Nd | Apoptosis | cCasp3 | CASP3 | Fluidigm | D3E9 | X | X | X |
| 143Nd | Apoptosis | cPARP | PARP1 | Fluidigm | F21-852 | X | X | |
| 144Nd | SUC | pan Akt | AKT1 | CST | 40D4 | X | X | X |
| 145Nd | Cell cycle | p-H3 [S28] | H3-4 | BioLegend | HTA28 | X | X | X |
| 146Nd | Cell cycle | Cyclin D1 | CCND1 | Abcam | SP4 | X | X | X |
| 147Sm | Stress/DNA damage | p-H2AX [S139] | H2AFX | Fluidigm | JBW301 | X | X | |
| 148Nd | Stress/DNA damage | p-Chk1 [S345] | CHEK1 | CST | 133D3 | | | X |
| 149Sm | TGFb pathway | p-Smad1 [S463/S465] / p-Smad8 [S465/S467] | SMAD1, SMAD9 | BD | N6-1233 | X | X | |
| 150Nd | Other | YAP | YY1AP1 | CST | D8H1X | X | X | X |
| 151Eu | MAPK pathway | p-MEK1/2 [S217/221] | MAP2K1, MAP2K2 | CST | 41G9 | X | X | |
| 152Sm | Akt/mTOR pathway | p-Akt [S473] | AKT1 | Fluidigm | D9E | X | X | |
| 153Eu | TGFb pathway | p-Smad2 [S465/467] / p-Smad3 [S423/425] | SMAD2, SMAD3 | CST | D27F4 | X | X | X |
| 154Sm | JAK/STAT pathway | p-Stat1 [T701] | STAT1 | BioLegend | A17012A | X | X | |
| 155Gd | TNFR pathway | p-NF-kB p65 [S536] | RELA | CST | 93H1 | X | X | |
| 156Gd | Stress/DNA damage | p-p38 [T180/Y182] | MAPK14 | Fluidigm | D3F9 | X | X | |
| 158Gd | JAK/STAT pathway | p-Stat3 [Y705] | STAT3 | Fluidigm | 4/P-STAT3 | X | X | |
| 159Tb | Stress/DNA damage | p-CDC25c [S216] | CDC25C | CST | 63F9 | X | X | |
| 160Gd | Cell cycle | Cyclin B1 | CCNB1 | BD | -11 GNS | X | X | X |
| 162Dy | Proliferation | Ki-67 | MKI67 | Fluidigm | B56 | X | X | X |
| 163Dy | Proliferation | p-RB [S807/S811] | RB1 | BD | J112-906 | X | X | X |
| 164Dy | TNFR pathway | IκBα | NFKBIA | Fluidigm | L35A5 | X | X | |
| 166Er | JAK/STAT pathway | CXCL1 | CXCL1 | R&D | 20326 | X | X | |
| 168Er | Akt/mTOR pathway | p-Akt [T308] | AKT1 | CST | D25E6 | X | X | |
| 169Tm | Other | GDF15 | GDF15 | Abcam | EPR19939 | X | | |
| 170Er | Akt/mTOR pathway | p-4EBP1 [T37/46] | EIF4EBP1 | CST | 236B4 | X | X | |
| 171Yb | MAPK pathway | pERK1/2 [T202/Y204] | MAPK3, MAPK1 | Fluidigm | D13.14.4E | X | X | |
| 172Yb | Stress/DNA damage | p-p53 [S15] | TP53 | CST | 16G8 | X | X | |
| 173Yb | Other | NICD | NOTCH1 | Abcam | ab8925 | X | | |
| 174Yb | SUC | total ERK1/2 | MAPK3, MAPK1 | CST | L34F12 | X | X | X |
| 175Lu | Akt/mTOR pathway | p-S6 [S235/236] | RPS6 | Fluidigm | N7-548 | X | X | |
| 176Yb | Other | Lamin B1 | LMNB1 | Abcam | EPR22165-121 | X | | X |

### S3. HNSCC data set

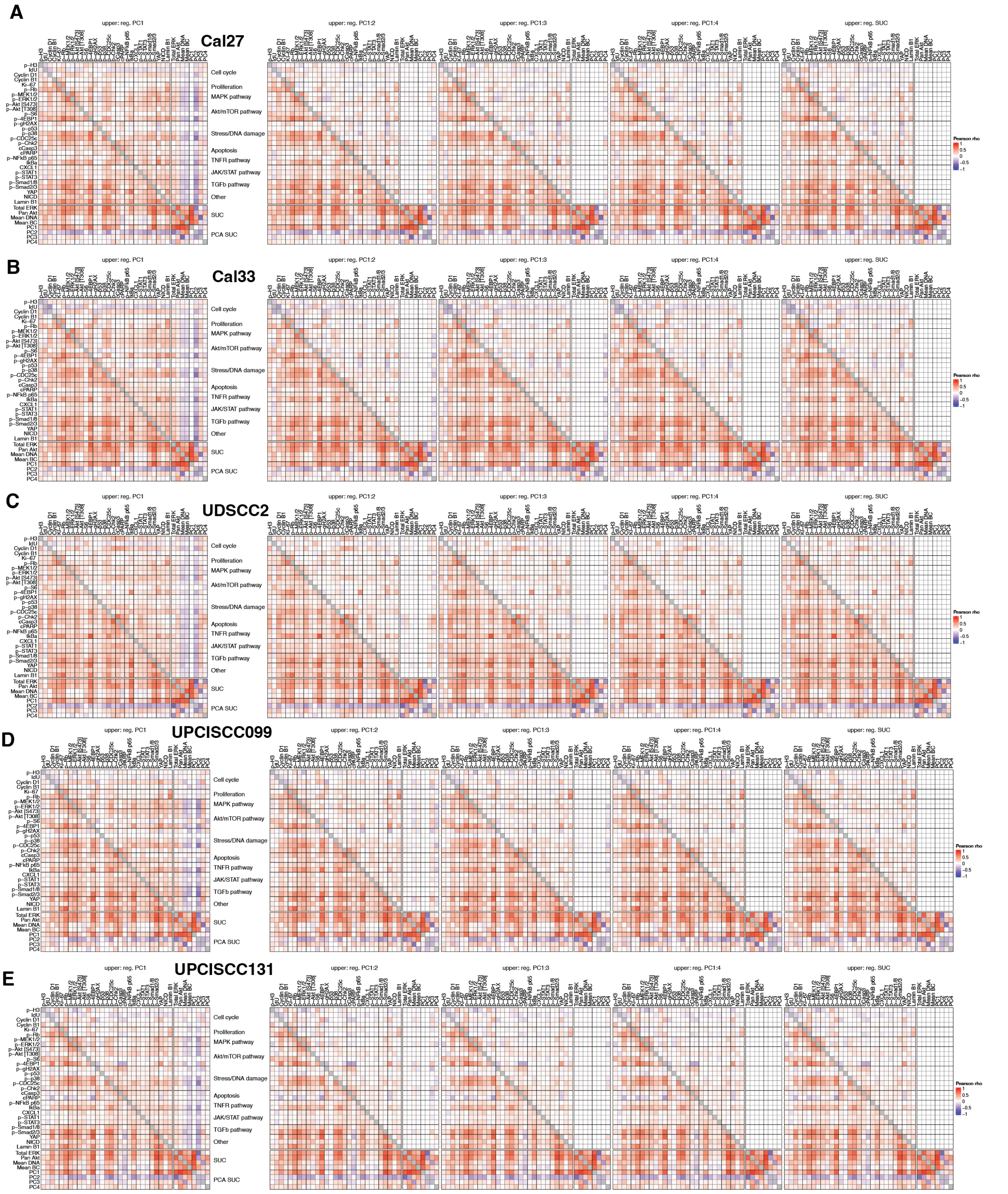

Figure S5: Correlation heatmap with the lower (upper) triangle showing the Pearson correlation coefficients between marker values before (after) RUCova based on PC1 (first) to PC1:4 (fourth) and the four SUCs (fifth), across cells from 0 and 10 Gy in the cell lines **A)** Cal27, **B)** Cal33, **C)** UDSCC2, **D)** UPSCISCC099, **E)** UPSCISCC131.

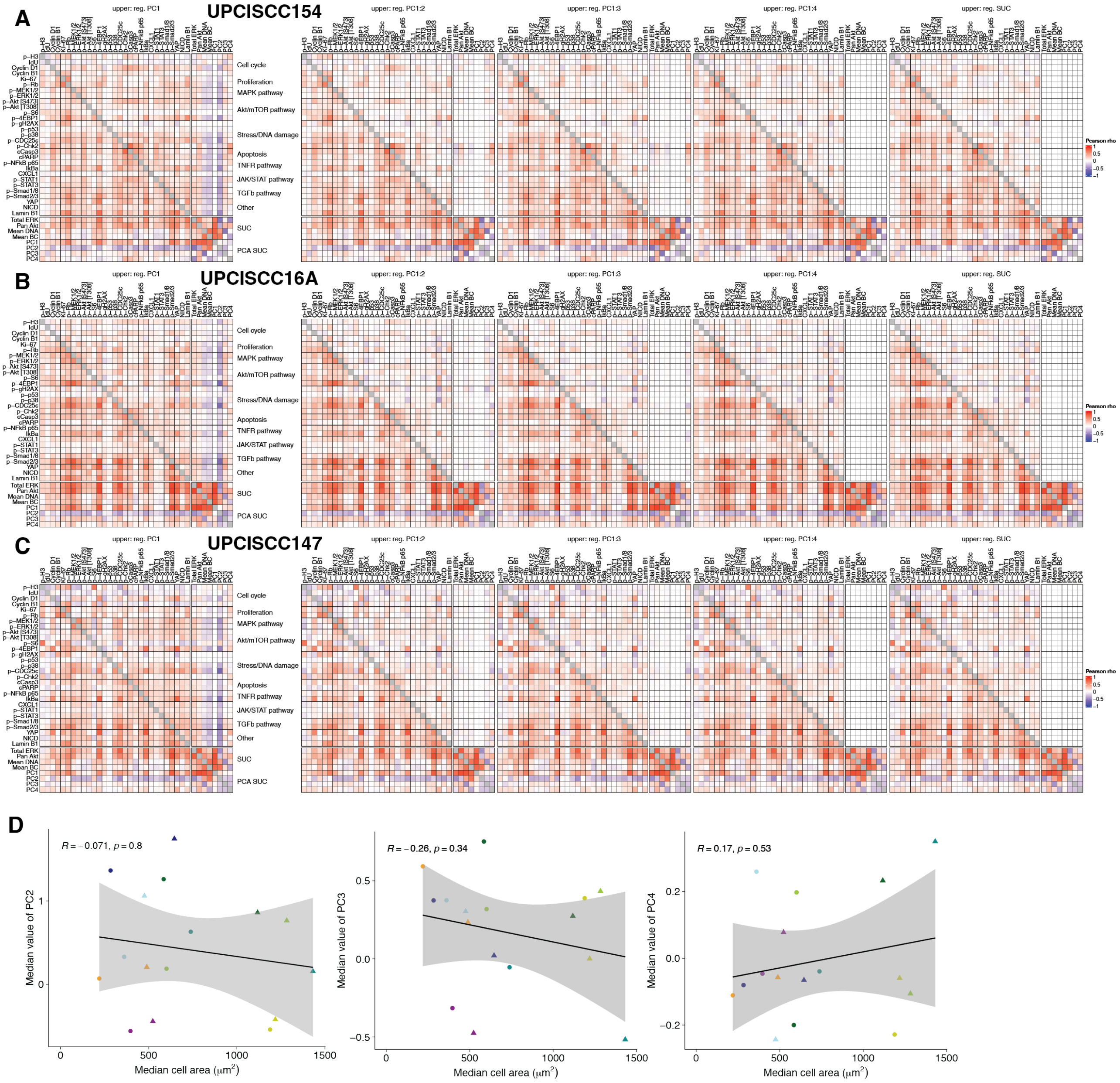

Figure S6: **A,B,C)** Correlation heatmap with the lower (upper) triangle showing the Pearson correlation coefficients between marker values before (after) RUCova based on PC1 (first) to PC1:4 (fourth) and the four SUCs (fifth), across cells from 0 and 10 Gy in the cell lines **A)** UPSCISCC154, **B)** UPSCISCC16A, **C)** UPSCISCC147. **D)** Scatter plots of measured mean cell area and mean value of PC2 (left), PC3 (middle), and PC4 (right).

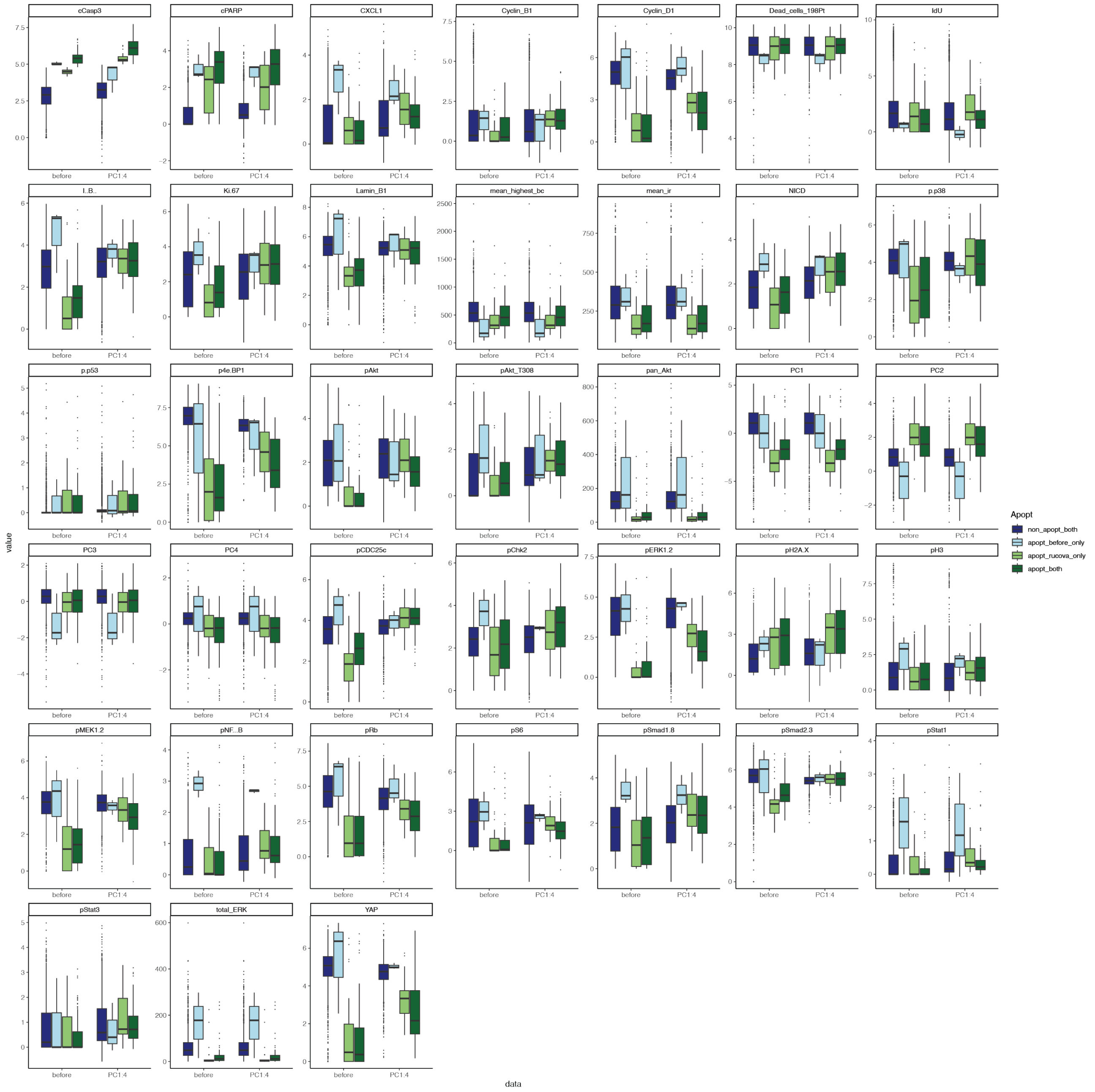

Figure S7: Boxplots of asinh-transformed marker expression irradiated UPCISCC131 cells according to their apoptotic status.

### S4. FACS-sorted data set

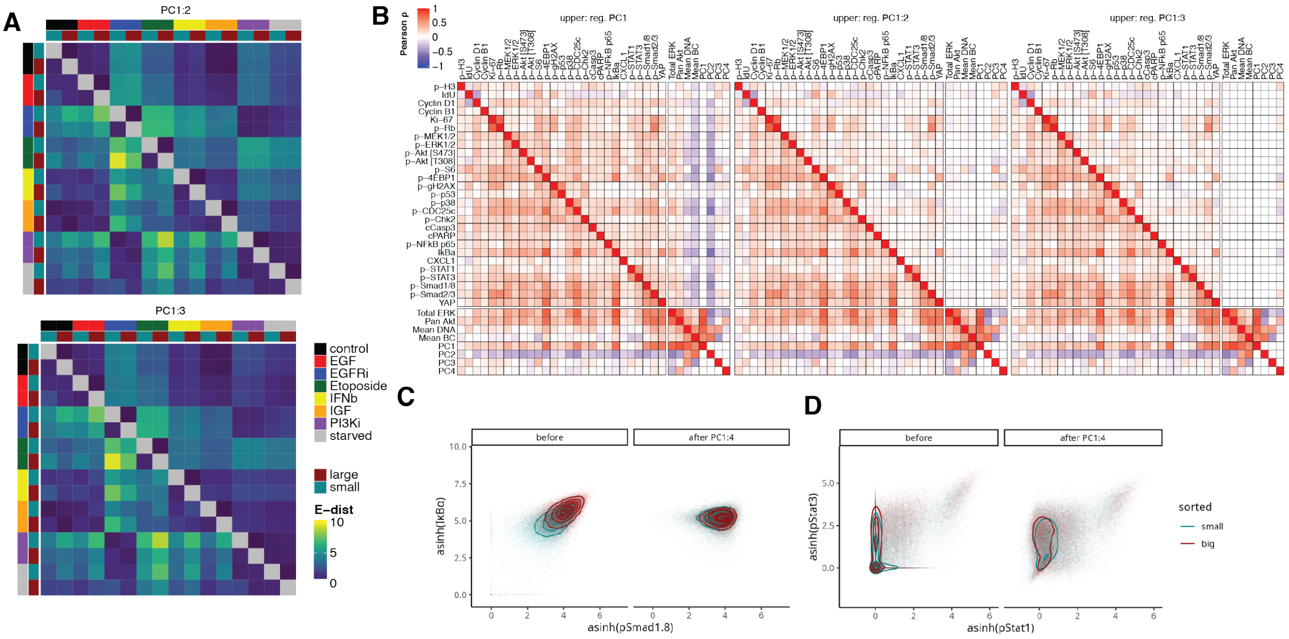

Figure S8: **A)** E-distance heatmap between conditions and sorted populations for data after RUCova using PC1 to PC2 (top) and PC1 to PC3 (bottom). **B)** Correlation heatmap with the upper triangle showing the Pearson correlation coefficients between marker values across all pertubations and sorted pupulations after RUCova based on PC1 (left), PC1 to PC2 (middle) and PC1 to PC3 (right). **C,D)** Scatter plots of Cal33 cells coloured by sorted population (small and large cells), before (left) and after (right) RUCova based on all four PCs. **C)** Asinh-transformed signals of p-Smad1/8 and I$\kappa$B$\alpha$. **D)** Asinh-transformed signals of p-Stat1 and p-Stat3.
